# RNA-binding protein Mub1 and the nuclear RNA exosome act to fine-tune environmental stress response

**DOI:** 10.1101/2020.07.01.181719

**Authors:** Adrien Birot, Cornelia Kilchert, Krzysztof Kus, Emily Priest, Ahmad Al Alwash, Alfredo Castello, Shabaz Mohammed, Lidia Vasiljeva

## Abstract

The nuclear RNA exosome plays a key role in mediating degradation and processing of multiple cellular RNAs. Recognition of the specific RNA substrates by the exosome is mediated by the RNA-binding co-factors. Transient binding of co-factors either to the exosome or the substrate RNAs as well as their rapid decay make identification of the co-factors challenging. Here, we employ a comparative poly(A)+ RNA interactome capture approach in the fission yeast exosome mutants to identify proteins that interact with poly(A)+ RNA in an exosome-dependent manner. Our analyses identified multiple proteins whose occupancy on RNA is altered in the exosome mutants including zinc-finger protein Mub1. Mub1 is required to maintain the levels of a subset of the exosome RNA substrates including mRNAs encoding for stress-responsive proteins. Removal of the zinc finger domain leads to loss of RNA suppression under non-stressed conditions, altered expression of heat shock genes in response to stress, and reduced growth at elevated temperature. These findings highlight importance of exosome-dependent mRNA degradation in buffering gene expression networks to mediate cellular adaptation to stress.

## INTRODUCTION

Regulation of RNA maturation and degradation is crucial to accurate gene expression (1). The nucleolytic RNA exosome complex has a central role in biogenesis of multiple types of transcripts produced by RNA polymerases I, II, and III (Pol I, II, and III) (2–9). The nuclear RNA exosome functions in RNA processing (3’end trimming) of stable non-coding (nc) RNA species such as ribosomal RNAs (rRNAs), transfer RNAs (tRNAs), telomerase RNA, small nuclear and nucleolar RNAs (snRNAs and snoRNAs) and quality control where it degrades precursors of unprocessed or incorrectly processed mRNAs and ncRNAs (2–11). The exosome also degrades unstable transcripts produced by Pol II (Cryptic Unstable Transcripts, CUTs) in yeast and multiple unstable ncRNAs such as promoter upstream transcripts (PROMPTs), enhancer RNAs and products of wide-spread premature transcription termination in higher eukaryotes (12–18). Recent studies have also demonstrated that the exosome not only removes unprocessed mRNAs but is also required for proper mRNA processing since exosome mutants show splicing and mRNA 3’end processing defects (19–23). Finally, the exosome regulates the levels of specific mRNA transcripts in response to environmental changes and is an important player in executing specific gene expression programmes during development (9, 23–27). In various human cell culture models, the activity of the exosome complex was shown to prevent cellular differentiation and maintain cells in an undifferentiated state by suppressing the expression of developmental regulators (28–31). Unsurprisingly, mutations in the nuclear exosome lead to severe neurodegenerative diseases in humans, such as spinal muscular atrophy and pontocerebellar hypoplasia (25, 27, 32).

The nuclear RNA exosome is a 3′–5′ exonuclease complex that consists of a 9-protein core (EXO-9) and two catalytic subunits, Rrp6 (EXOSC10) and Dis3/Rrp44 (hDIS3). EXO-9 forms a double-layered barrel-like structure that comprises six ribonuclease (RNase) (PH)-like proteins (Rrp41, Rrp42, Rrp43, Rrp45, Rrp46, and Mtr3) and three S1/K homology (KH) “cap” proteins (Rrp4, Rrp40, and Csl4) (33). The two catalytic subunits occupy opposite ends of EXO-9 to constitute EXO-11 (34, 35). Rrp6 is located at the top of the S1/KH cap ring near the RNA entry into the channel formed by the exosome cap and core, and Dis3 is found at the bottom of EXO-9 channel near the RNA exit pore. Both Rrp6 and Dis3 are 3′–5′ exonucleases, but the latter also has endonucleolytic activity (2). In yeast, Rrp6 is restricted to the nucleus and Dis3 is found in both nuclear and cytoplasmic compartment (36, 37).

The conserved helicase Mtr4 is essential for RNA degradation by the exosome (38). In the fission yeast, *Schizosaccharomyces pombe* (*S. pombe*), Mtr4 shares its function with the highly homologous Mtr4-like helicase (Mtl1) (39). Mtr4/Mtl1 interacts with RNA-binding proteins and the exosome and was proposed to play a central role in exosome recruitment to substrate RNAs (1, 40–42). In addition to the exosome core, Mtr4/Mtl1 co-purify with RNA-binding proteins involved in substrate recognition. In *Saccharomyces cerevisiae* (*S. cerevisiae*), Mtr4 is a part of the TRAMP complex (Trf4/5–Air1/2–Mtr4) which consists of two Zinc-finger proteins Air1 and Air2 (only Air1 in fission yeast), poly(A) polymerases Trf4 and Trf5 and Mtr4 (43–45). The TRAMP complex is recruited to RNA by the RNA and Pol II-binding protein Nrd1 during transcription (14, 46–49). However, in contrast to *S. cerevisia*e, the TRAMP complex seems more specialized in regulation of rRNA processing in fission yeast and human (38).

In fission yeast, Mtl1 interacts with the conserved YTH-domain containing protein Mmi1 and associated proteins, Red1, Iss10 and Erh1 (16, 39, 50–52). Mmi1 is needed for degradation of a selected group of mRNAs encoding for proteins involved in meiosis, cell cycle regulation, biosynthetic enzymes, RNA biology and ncRNAs by the exosome. Mmi1 is co-transcriptionally recruited to these transcripts by binding to an UUAAAC sequence motifs known as ‘determinants of selective removal’ (DSRs), and this is proposed to lead to their degradation (23, 24, 26, 53). It was recently demonstrated that Mmi1 interaction with Red1 and Mtl1 as a part of MTREC/NURS (Mtl1-Red1 Core / Nuclear RNA Silencing) complex is important to mediate recruitment of the exosome to RNAs (54). Mmi1 is also important for mRNA quality control and in particular for degradation of inefficiently spliced mRNAs and proper transcription termination of selected transcripts (19, 23, 55). In addition to Mmi1, Iss10, Erh1 and Red1, Mtl1 also co-purifies with other factors that have been functionally linked to the exosome regulation: the Zinc-finger protein Red5, the poly(A) binding protein Pab2, RRM (RNA-Recognition-Motif) and PWI (Pro-Trp-Ile signature) domain containing protein Rmn1 and the CAP-binding proteins Cbc1, Cbc2 and Ars2 (16, 39). Nevertheless, the mechanisms by which these factors regulate substrate recognition and exosome targeting to substrate RNAs remain poorly understood.

In addition to a group of RNAs recognised by Mmi1, levels of multiple other transcripts have been reported to increase in nuclear exosome mutants suggesting that additional Mmi1-independent mechanisms could contribute to their recognition (16, 39). We therefore employed a quantitative proteomics approach to identify RNA-binding proteins that could potentially be involved in mediating exosome targeting to RNAs. We hypothesized that the association with RNA of potential exosome co-factors should increase upon stabilisation of the substrate RNAs in the exosome mutants. We compared the poly(A)+ transcriptomes and poly(A)+ RNA-bound proteomes from control cells and three different exosome mutants (*mtl1-1, rrp6Δ* and *dis3-54*). Our data suggest that the nuclear exosome plays a more prominent role in controlling the fission yeast transcriptome than previously anticipated. In addition, analysis of the impact of the mutations in different exosome subunits on poly(A)+ bound proteome has identified proteins with increased RNA binding that could function as potential regulators of the exosome in fission yeast. We selected ten RNA-binding proteins with significantly altered RNA association and demonstrate that deletion of each of these proteins phenocopies at least one of various phenotypes typical of inactivated nuclear exosome, namely suppression of transposon RNAs, telomeric silencing, and nuclear RNA retention. We focus on the uncharacterised zf-MYND (MYeloid, Nervy, and DEAF-1) protein Mub1, which is highly enriched on poly(A)+ RNA in exosome mutants. Mub1 physically interacts with the exosome and its deletion leads to increased levels of a specific subset of exosome substrates supporting its role in exosome regulation.

## RESULTS

### Poly(A)+ RNA interactome capture in the exosome mutants

We had previously employed an unbiased quantitative proteomic approach, RNA interactome capture (RIC), to assess how mutation of various exosome components affects association of the entire complex with polyadenylated RNA (poly(A)+) (56). Specifically, we analysed: *rrp6Δ*, lacking exonuclease Rrp6; *dis3-54*, a Dis3 mutant which contains an amino acid substitution (Pro509 to Leu509) located within RNB domain that was reported to show reduced catalytic activity (57, 58) and *mtl1-1*, a mutant of the helicase Mtl1, which has mutations in the region surrounding the arch domain (39) (Figure 1A). All three mutants are defective in RNA turnover and accumulate known targets of the exosome (19, 39, 59). To gain further insights into the regulation of the exosome complex, we re-analyzed the RIC data to identify RBPs that show increased interactions with poly(A)+ RNA in these mutants. The underlying hypothesis behind this approach was that stabilization of RNAs targeted by the exosome would facilitate the capture of proteins that are enriched on these RNAs, and that are likely to be functionally linked to the exosome (Figure 1B). The three mutant strains were cultured alongside a WT control in the presence of 4-thiouracil (4sU) to facilitate RNA-protein crosslinking with 365nm UV light. Following UV crosslinking, poly(A)+ RNA was enriched by oligo-d(T) selection and poly(A)+ RNA-associated proteins were identified by mass spectrometry (Figure 1B). The abundance of individual proteins in the whole cell extract (WCE) was used to normalise the RIC data (see Material and Methods) as in (56, 60). The RIC/WCE ratio was used to determine the enrichment of each individual protein on poly(A)+ RNA in the mutants relative to the WT (Supplementary Table 1). Additionally, RNA sequencing (RNA-seq) was carried out for oligo-d(T) enriched samples to assess levels of individual polyadenylated RNAs in the WT cells and exosome mutants (Figure 1B).

**Figure 1:**
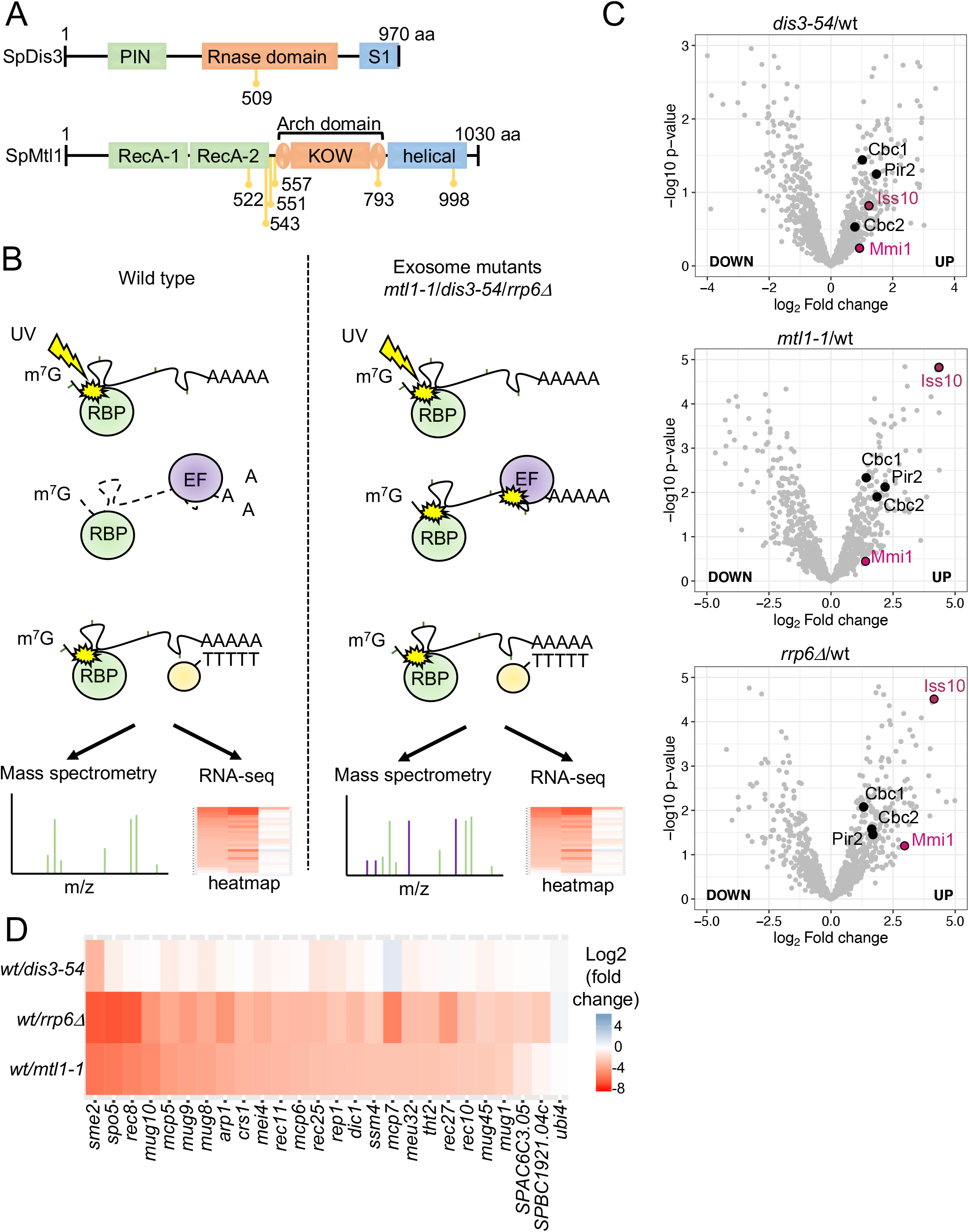
Comparative poly(A)+ RNA interactome capture from fission yeast exosome mutants. **A**. Schematic diagram describing domain organisation of *S. pombe* (Sp) Dis3 and Mtl1 and position of the mutations in *dis3-54* (P509L) and *mtl1-1* (I522M, L543P, Y551H, L557P, D793G, A998V) shown in yellow. **B**. Schematic diagram describing the poly(A)+ interactome capture approach. Cells were grown in the presence of 4-thiouracil (4sU) and exposed to UV (3J/cm^2^) to allow protein-RNA crosslinking. Poly(A)+ RNA and associated proteins were pulled down by oligo-d(T) beads and subjected to RNA sequencing and mass spectrometry analyses (RBP= RNA-Binding Proteins, EF = Exosome Factors, involved in recognition of the exosome substrates such as Mmi1). **C**. Volcano plots showing distribution of Mmi1, Iss10, Cbc1, Cbc2 and Ars2 (Pir2 in *S. pombe*) enrichment on poly(A)+ RNA in the exosome mutants. In the volcano plot, *p*-values (-log10, moderated Student’s *t-test*) are plotted against the ratio of log2-fold changes in mass spectrometry (MS) intensities for the whole-cell extract normalised proteomes of mutant versus WT cells recovered from the oligo-d(T) pull-downs of UV-crosslinked samples (3 J/cm^2^). In all panels, individual proteins are depicted as a single dot. **D**. Heat map analysis of poly(A)+ RNA-seq showing differential expression of the Mmi1 regulon in exosome mutants compared to WT.

### Exosome target RNAs are enriched in the poly(A)+ RIC samples of the exosome

First, to confirm that exosome target RNAs were indeed overrepresented in the RIC samples of the different exosome mutants, we analysed them by RNA-seq. Consistent with the function of the exosome in degradation of ncRNAs (snRNAs, snoRNAs, CUTs), levels of multiple nc transcripts were increased in all three exosome mutants (Figure EV (Expanded View)1A, Supplementary Table 3). In agreement with previously reports, a high number of mRNAs (835, 1078 and 317 for *mtl1-1, rrp6Δ* and *dis3-54* respectively, >1.5-fold, *p*-value<0.05) is also increased in the exosome mutants confirming that the nuclear exosome regulates multiple mRNAs in addition to its well-described role in degradation of ncRNAs (Figure EV1B-D) (16, 39). The majority of the transcripts increased in *rrp6Δ* were also increased in *mtl1-1* (~80%) in agreement with the suggested functional connection between Mtl1 and Rrp6 (39, 54). Compared to *rrp6Δ* and *mtl1-1* mutants, a lower number of RNAs are increased in *dis3-54* mutant (1373, 1718 and 623 for *mtl1-1, rrp6Δ* and *dis3-54* respectively, >1.5-fold, *p*-value<0.05), possibly reflecting Dis3 is only partially inhibited by these mutations under conditions tested (Figure EV1A). However, approximately half of the transcripts that show increased levels in the Dis3 mutant (54%) were also dependent on Mtl1. The reason for the observed differences in behaviour of the two mutants is likely because the single P509L mutation in *dis3-54* may cause a milder effect on Dis3 function when compared to the multiple mutations present in *mtl1-1* (Figure 1A). Our data supports a model where Mtl1 regulates both exosomes-associated nucleases (Rrp6 and Dis3), with Rrp6 being perhaps slightly more dependent on Mtl1 than on Dis3, and confirms that exosome target RNAs are enriched in the RIC samples.

### Effect of the mutations in the exosome subunits on poly(A)+ RNA interactome

To assess how inactivation of the different subunits of the exosome impact RNA-protein binding profile, we next performed comparative analysis of proteins differentially enriched in interactomes from *mtl1-1, rrp6Δ* and *dis3-54* compared to WT. For each RIC experiment, proteins that are detected at least in two out of three biological repeats are used in the analysis (see Material and Methods), resulting in a total of total of 1146 proteins. Interestingly, 152, 180 and 83 proteins are 2-fold enriched in *mtl1-1, rrp6Δ* and *dis3-54* lines over WT, respectively (*p*-value<0.1, Figure EV1E and Supplementary Table 2). Consistent with the RNA-seq data, the overlap between the list of RBPs stimulated in *rrp6Δ* and *mtl1-1* cells is substantially larger than in *dis3-54* and *mtl1-1*, further supporting a functional link between Mtl1 and Rrp6 (Figure EV1E). On the other hand, we detected 137, 95 and 86 proteins whose abundance on poly(A)+ RNA was reduced in *mtl1-1, rrp6Δ* and *dis3-54* (*p*-value<0.1, Figure EV1F and Supplementary Table 2). Next, we performed gene ontology (GO) analyses across the set of RBPs differentially regulated in the three different mutants to assess whether they are linked to a specific biological process. The analysis revealed that nuclear proteins are most noticeably affected in all three exosomes mutants including *dis3-54* mutant, even if both nuclear and cytoplasmic forms of the exosome are compromised in this strain (Figure EV1G, Supplementary Table 4). This agrees with the prominent role of the exosome in nuclear RNA metabolism. Additionally, all mutants exhibit alterations of RBPs involved in “mRNA metabolic process” (GO:0016071) and “ribosome biogenesis” (GO:0042254) (Supplementary Table 4), suggesting a profound reorganisation of RNPs in the exosome mutants. In contrast, the GO term “cytoplasmic translation” (GO:0002181) was significantly over-represented among the RBPs with reduced RNA-binding activity in the catalytic exosome mutants. Indeed, “cytoplasmic translation” annotated proteins compress 22 out of 95 downregulated RBPs in *rrp6Δ* (*p*-value = 2,02^e-7^) and 31 out of 86 in *dis3-54* (*p*-value = 7,16^e-17^). This may reflect a change in subcellular localisation of poly(A)+ RNA in these strains. Indeed, it was previously reported that poly(A)+ RNA is retained in the nucleus in Rrp6 mutants (61–63).

To assess if the exosome mutants exhibit a defect poly(A)+ RNA localisation, we performed RNA FISH with oligo(dT) probes. Notably, *rrp6Δ* and *mtl1-1* mutants showed accumulation of poly(A)+ RNA inside the nucleus (Figure EV1H-I). Consistent with the *dis3-54* milder phenotypes (see above), this line shows modest poly(A)+ RNA accumulation in the nucleus (Figure EV1J). Milder nuclear retention of poly(A)+ RNA observed in *dis3-54* compared to other exosome mutants might be due to only partial inactivation of Dis3 in this mutant as discussed above. However, this might also indicate that Dis3 plays less dominant role in regulation of RNA in the nucleus compared to Mtl1 and Rrp6. Interestingly, even though the accumulation of poly(A)+ RNA is only mild in *dis3-54*, nuclear proteins are strongly overrepresented among proteins with increased RNA binding in this mutant (Figure EV1G), suggesting that the exosome has more a prominent impact on nuclear rather than cytoplasmic RNPs.

Altogether, RIC analyses of exosome mutants demonstrate significant changes in protein-RNA interactions in exosome-deficient cells, highlighting the important role of the exosome in nuclear RNP metabolism.

### Comparative RIC identifies known RNA-binding proteins linked to the nuclear RNA exosome

To validate that the comparative interactome capture approach leads to enrichment of exosome co-factors, we first analysed how RNA occupancy of Mmi1 is affected in the exosome mutants as a proof of principle. As predicted, Mmi1 association with poly(A)+ RNA is 1.5, 2 and 4 times increased in *dis3-54, mtl1-1* and *rrp6Δ* RIC, respectively, compared to WT cells (*p*-value=0.768, *p*-value=0.039 and *p*-value=0.001) (Figure 1C). The increased RNA association of Mmi1 correlates with the increased abundance of Mmi1 RNA substrates in the exosome mutants (Figure 1D). Indeed, well-characterised Mmi1 mRNA targets such as transcripts encoding for meiotic proteins are noticeably enriched in the *rrp6Δ* and *mtl1-1* mutants and to some extend in *dis3-54*. Moreover, association of other known exosome co-factors with poly(A)+ RNA is also increased in the exosome mutants. For example, the RNA-binding activity of Iss10 protein, which is proposed to interact with Mmi1, is increased from 2 to more than 10-fold in *dis3-54* and *mtl1-1, rrp6Δ* RIC respectively over WT cells (p-value=0.15, p-value=3.26×10-5 and p-value=1.73×10-5). Similarly, the RNA-binding activity of the Cap-binding complex Cbc1, Cbc2 and Ars2 is also increased (>4 fold) in the exosome mutants (Supplementary table 1). Altogether, these results illustrate the capacity of RIC to discover RBPs with bona fide roles in exosome targeting to substrate RNAs.

### Identification of RBPs with functional links to the exosome

To uncover cellular RBPs with novel roles in the exosome function, we selected proteins containing classical RNA-binding domains (64) out of the list of proteins whose RNA-binding activity is increased (>2 fold) in the exosome mutants. From these RBPs, we focused on proteins annotated as “uncharacterised”. This resulted in a list of 10 proteins (Figure 2A). Next, the candidate RBPs were deleted to test whether they recapitulate phenotypes associated with compromised exosome function. First, we assessed whether nuclear retention of poly(A)+ RNA, characteristic for the exosome mutants, is observed in the deletion strains using poly(A)+ FISH (Figure 2B). *rrp6Δ* strain shows strong nuclear retention phenotype for poly(A)+ RNA that results in 2 fold higher nuclear/cytoplasmic ratio compared to WT (Figure 2C). As a result of the analyses of subcellular distribution of poly(A)+ RNA across all strains tested, we assigned the 10 candidates to 3 different groups according to the ratio of poly(A)+ signal between the nucleus and the cytoplasm, which ranges from moderately increased to no change (Figure 2C and Figure EV2A). The deletion of *SPCC126*.*11c* leads to noticeable nuclear retention of poly(A)+ RNA (ratio of poly(A)+ signal ~1.5). A mild retention of poly(A)+ RNA in the nucleus is observed in strains lacking Srp40, Pof8, SPAC222.08, SPBC530.08, SPBC16G5.16, Swt1 and Mlo1 with a ratio around 1.2 (Figure 2C and Figure EV2A). In contrast, no retention is observed upon deletion of *SPAC31F10*.*10c* or *SPABC18H10*.*09*.

**Figure 2:**
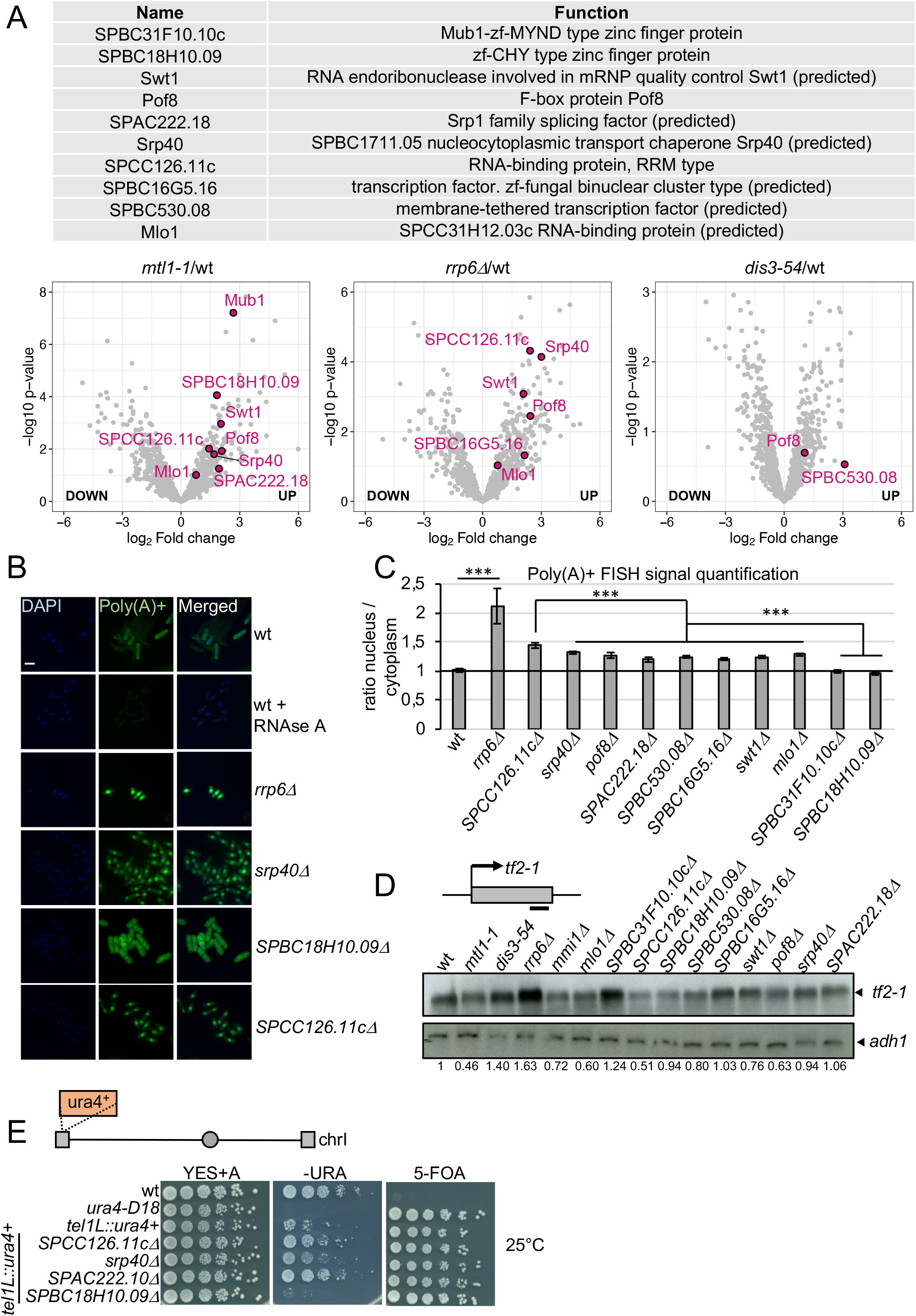
Comparative RIC uncovers putative novel exosome factors. **A**. List of the candidate proteins enriched on poly(A)+ RNA in the exosome mutants, their predicted function and volcano plot showing their distribution in *mtl1-1, rrp6Δ* and *dis3-54* mutants. **B**. Poly(A)+ RNA FISH illustrating cellular localisation of poly(A)+ RNA in the indicated strains. DAPI is shown in blue. Poly(A)+ RNA is visualised in green. Scale bar = 10µm. **C**. Quantification of Poly(A)+ RNA FISH analysis (in Figures 2B and EV2A), (***: *p*-value<0.0001, un-paired *t-test*). **D**. Analyses of *tf2-1* mRNA levels by northern blot in the indicated strains. Band corresponding to *tf2-1* is indicated with an arrow. Schematics shows location of the probe with a black bar. The *adh1* mRNA is used as a loading control. Numbers indicate the relative abundance of *tf2-1* transcript normalised to *adh1* levels. **E**. Serial dilutions of the indicated strains plated on complete medium supplemented with adenine (YES+A), medium lacking uracil (-URA) and containing 5-FOA. Cells are grown at 25°C. A schematic representation of the Chromosome I (Chr I) depicting centromeric and telomeric regions (grey circle and grey rectangles correspondently). Position of *ura4*^*+*^ reporter is shown.

Next, we assessed the steady-state levels of RNAs that are up-regulated in the exosome mutants but are independent of Mmi1, such as the RNA produced from the Tf2 retro-transposable element (*SPAC9*.*04*). Increase in *tf2-1* RNA levels is only observed in *SPBC31F10*.*10cΔ* with a ratio of 1.24 compared to WT, suggesting that SPBC31F10.10c controls levels of this RNA, similar to Rrp6 (Figure 2D).

Exosome mutants also show defects in heterochromatic silencing (39, 65). This results in accumulation of heterogeneous RNAs produced from telomeric and centromeric regions of the *S. pombe* genome, although the underlying mechanism remains unclear. To test whether the candidate proteins might contribute to the repression of heterochromatic transcripts, each individual deletion mutant was crossed with a strain bearing a *ura4+* reporter gene inserted within the transcriptionally silent telomeric region of chromosome I. We then monitored the ability of these reporter strains to grow on - URA plates or plates containing 5-Fluoroorotic acid (5-FOA), a compound that is toxic when a functional version of the *ura4* gene is expressed, at 25°C and 30°C. The deletion of either *SPCC126*.*11c, srp40* or *SPAC222*.*18* led to moderate growth on –URA plates and attenuated growth on 5-FOA compared to WT (Figure 2E and Figure EV2B), suggesting that each of these candidates might play a role in heterochromatin formation or maintenance.

### Mub1 regulates levels of stress-induced mRNAs targeted by the exosome

We decided to investigate *SPBC31F10*.*10c* further, because its deletion led to increased levels of *tf2-1* RNA targeted by the exosome. Additionally, Mub1 shows a >6-fold increase in *mtl1-1* RIC (*p*-value= 5,41E-08) (Figure 2A). *SPBC31F10*.*10c* is the homologue of *S. cerevisiae* Mub1 (multi-budding 1), and contains an Armadillo-type domain and a potential nucleic acid binding region represented by a zf-MYND domain. To assess the contribution of Mub1 in regulation of RNA levels we carried out transcriptome analyses by RNA-seq in WT and mutant strains: *mub1Δ, mtl1-1, mtl1-1 mub1Δ*. RNA-seq data was normalised to *S. cerevisiae* spike-in for calibration (see Material and Methods). 248 transcripts (162 mRNAs and 86 ncRNAs) were upregulated more than 1.5-fold (*p*-value<0.05) in *mub1Δ* cells (Figure 3A, Supplementary Tables 5-6). The GO analysis of mRNAs increased in *mub1Δ* cells showed a significant enrichment for GO “Core Environmental Stress Response induced (CESR)” (46.25% (74/160), *p*-value = 1.55246^e-28^, AnGeLi tool (66)) implying that Mub1 may help to fine-tune the cellular response to stress. 144 transcripts (92 mRNAs and 52 ncRNAs) that are affected by Mub1 deletion were also up-regulated in *mtl1-1* (Figure 3A-B), suggesting that Mtl1 and Mub1 act in the same pathway. The GO term “Core Environmental Stress Response induced” was even more strongly enriched among mRNAs upregulated in both *mtl1-1* and *mub1*Δ (64.44% (58/90), *p*-value = 3.32315^e-32^, AnGeLi tool (66)), potentially indicating a role for Mub1 and Mtl1 in repressing the stress response under non-stress conditions. To validate the RNA-seq data, increased steady-state levels of *gst2* and *SPCC663*.*08c* mRNAs in *mub1Δ* and *mtl1-1* mutants were confirmed by northern blot analyses (Figure 3B). No additive effect was observed in the double mutant *mtl1-1 mub1Δ* compared to the single mutants, which is consistent with Mtl1 and Mub1 acting in the same pathway (Figure 3B). To assess whether Mub1 and the exosome physically interact, we carried out co-immunoprecipitation experiments. Mub1 co-purified with the Rrp6 subunit of the exosome, supporting a physical link between Mub1 and the nuclear exosome (Figure 3C).

**Figure 3:**
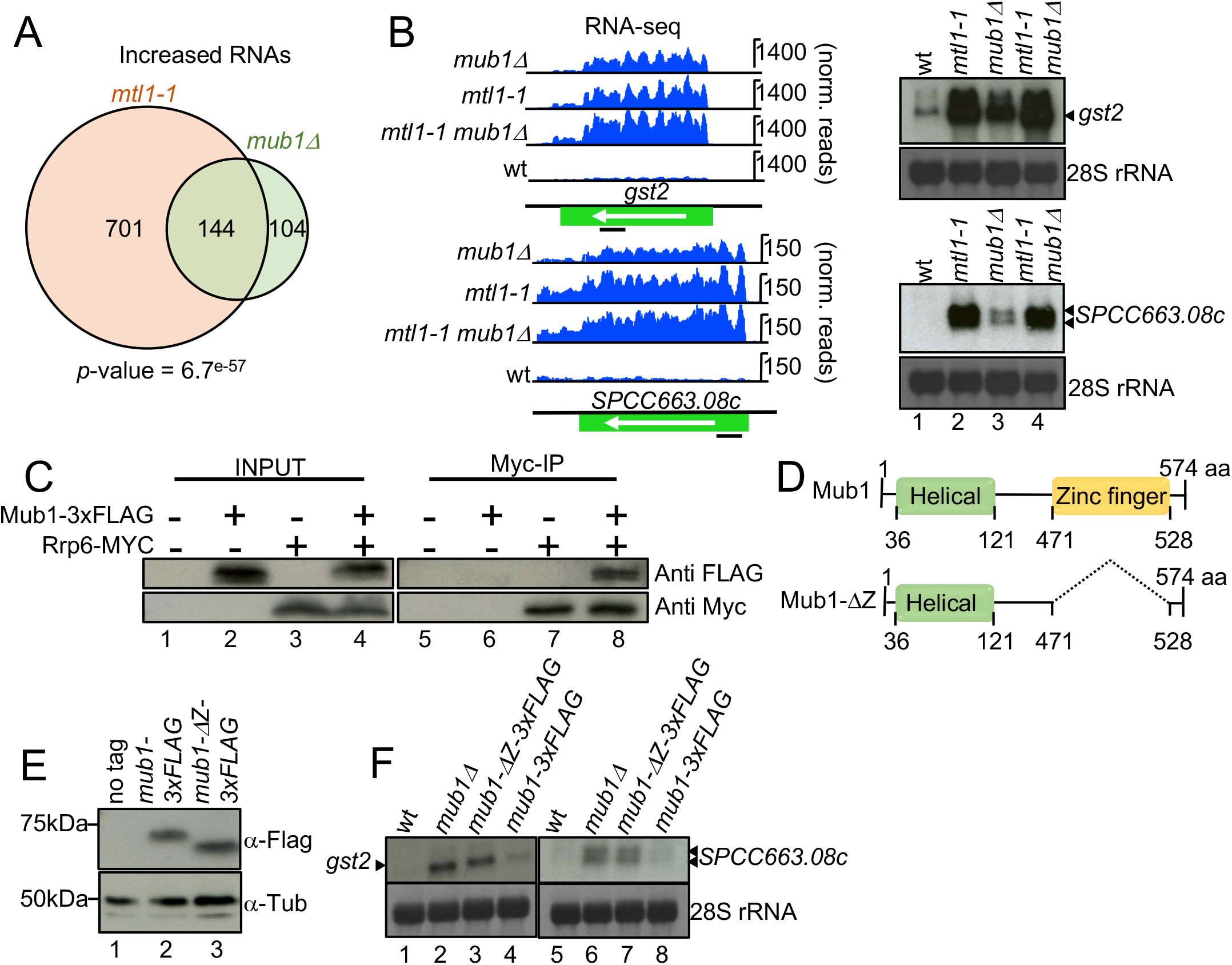
Mub1 protein functions in maintaining levels of the exosome RNA targets. **A**. Venn diagram showing overlap between RNAs that show increased levels in *mtl1-1* and *mub1Δ* (>1.5-fold, *p*-value<0.05) **B**. Genome browser snapshots showing RNA-seq data and northern blot analyses for two representative transcripts (*gst2* and *spcc663*.*08c*) on total RNA isolated from *mtl1-1, mub1Δ* and WT cells. Bands corresponding to *gst2* and *spcc663*.*08c* are indicated with arrows. Positions of the probes are indicated with black bars. 28S ribosomal RNAs (rRNA) visualised with methylene blue served as loading control. **C**. Co-immunoprecipitation of Rrp6-Myc with Mub1-3xFLAG. **D**. Graphical representation of WT Mub1 and Mub1-*Δ*-Zinc-Finger domain. **E**. Western blot showing levels of endogenously expressed Mub1-3xFLAG and Mub1-*Δ*Z-3xFLAG proteins. **F**. Analyses of *SPCC663*.*08c* and *gst2* mRNAs from *mub1-3xFLAG* and *mub1-ΔZ-3xFLAG* strains by northern blot as in **B**.

### The MYND zinc finger domain is required for regulation of RNA levels

The strong enrichment of Mub1 in the poly(A)+ RNA interactome suggested that Mub1 is an RBP. Mub1 contains a predicted zf-MYND domain (471-528), which could mediate its interaction with RNA. Like Mub1, multiple factors associated with the exosome contain Zinc-finger domains. In fission yeast these are: Red1, Red5 and Air1, a component of the TRAMP complex (51, 67, 68). The zinc-finger domains of Red1 and Red5 were shown to be important for RNA degradation by the exosome as mutations in this region lead to RNA stabilisation (51, 68). To test whether the zf-MYND domain is required for Mub1 function, we deleted the region between amino acids 471 and 528 (Figure 3D). Deletion of the zf-MYND domain led to the expected change in the size of the protein, which was assessed by visualising FLAG-tagged protein by Western blotting (Figure 3E). Addition of a triple FLAG tag to full-length Mub1 did not have any effect on cellular protein levels or cell growth, nor did it induce the spheroid cell shape typical of a *mub1* deletion (69) (Figure EV3A-C). The protein level of Mub1*Δ*471-528-3xFLAG (Mub1-*Δ*Z-3xFLAG) was comparable to Mub1-3xFLAG, suggesting that the deletion did not affect the stability of the truncated protein (Figure 3E). To test the importance of the zinc-finger domain for regulating mRNA levels, we assessed levels of *gst2* and *SPCC663*.*08c* mRNAs in *mub1Δ471-528-3xFLAG* mutants by northern blot. Like in *mub1Δ*, levels of both mRNAs were increased in the *mub1Δ471-528-3xFLAG* mutant (Figure 3F). In addition, cells expressing the truncated Mub1*Δ*471-528-3xFLAG displayed the characteristic rounded shape of *mub1Δ* cells (data not shown). Taken together, these results suggest that the zinc-finger domain of Mub1 is essential for its function.

### Mub1 plays a role in the heat shock response

We then investigated the physiological role of Mub1. We noticed that growth at 37°C is impaired in *mub1Δ* and Mub1-*Δ*Z-3xFLAG, which are both deficient in the regulation of mRNA levels (Figure 4A). Heat shock triggers the expression of genes involved in the core environmental stress response, many of which are dysregulated in *mub1Δ* and *mtl1-1* (see above), including *hsp16*, which encodes a heat shock protein (70) (Figure 4B, compare lanes 2-4 to 1). To test whether this dysregulation extended to stress conditions, we assayed, *hsp16* mRNA levels after induction of the heat shock response (4 hours at 37°C). As expected, levels of *hsp16* mRNA were induced after heat shock in WT cells, but the heat shock-dependent induction was amplified in *mub1Δ* and *mtl1-1*, or the double mutant (compare lanes 6-7 to lane 5). Our data highlight a role of Mub1 and the exosome complex in suppressing the heat shock response potentially through increased turnover of stress-responsive mRNAs. A reverse mechanism was recently described in budding yeast, where the Rrp6 component of the exosome was found to be required for the full induction of cell wall integrity (CWI) genes, thereby promoting cell survival during heat shock (71). Similarly, the exosome was shown to be recruited to heat shock genes in heat-stressed *Drosophila* Kc cells (72). This suggests that a function of the exosome in modulating environmental adaptation is conserved and can act on multiple levels. Further works will assess by which mechanism Mub1 contributes to this regulation.

**Figure 4:**
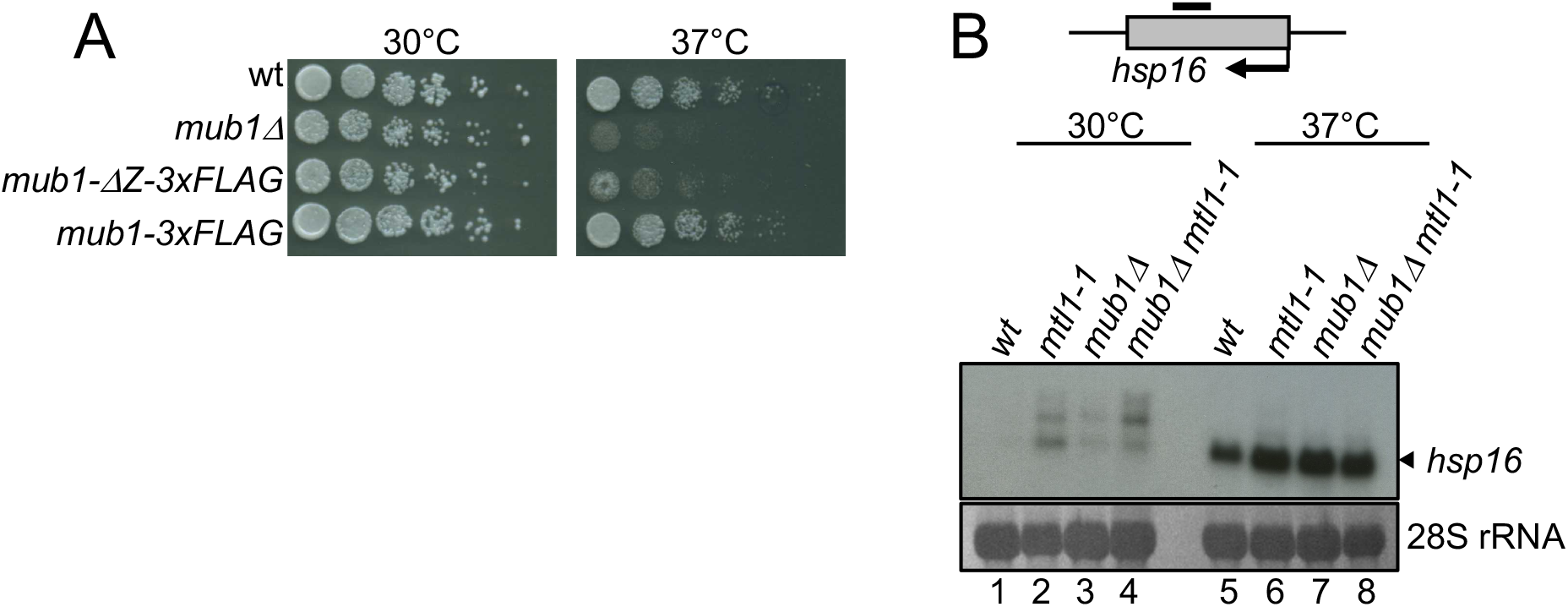
Mub1 and nuclear exosome are involved in regulation of heat shock response. **A**. Serial dilutions of the indicated strains plated on complete medium (YES+A) at the indicated temperatures. **B**. Analyses of *hsp16* mRNAs levels by northern blot using total RNA isolated from the indicated strains grown at the indicated temperatures. Band corresponding to *hsp16* mRNA is indicated with arrow. Schematic diagram shows location of the probe with a black bar. 28S ribosomal RNAs (rRNA) visualised with methylene blue serves as loading control.

Taken together, our RIC analysis identified several potential exosome regulators, including Mub1, SPCC126.11c, Srp40 or SPAC222.18, whose deletion phenocopies key aspects of exosome dysfunction, suggesting a relationship between these factors and the nuclear RNA exosome. Besides Mub1, which we characterize in detail, SPCC126.11c and Srp40 were previously reported to co-purify with the exosome, suggesting a direct functional link between these proteins and the exosome (50, 73). Moreover, a recent study reports that SRSF3, the human orthologue of SPAC222.18, is involved in exosome recruitment via the Nuclear Exosome Targeting (NEXT) complex, raising the possibility that the role of the serine rich protein SPAC222.18 is conserved in *S. pombe (74)*. Further studies are needed to test whether this is indeed the case.

## MATERIAL AND METHODS

### Yeast strains and manipulations

General fission yeast protocols and media are described in (75). All strains are listed in Supplementary Table 7. Experiments were carried out using YES medium at 30°C unless stated otherwise. Gene deletions and epitope tagging were performed by homologous recombination using polymerase chain reaction (PCR) products (76). All oligos are listed in Supplementary Table 8. Protein extracts and western blotting were done as described in (77).

### Co-Immunoprecipitation

Total-protein extracts were prepared from cycling log-phase cells. Cells were collected, rinsed in ice-cold phosphate-buffered saline (PBS)-1 mM phenylmethylsulfonyl fluoride (PMSF), and frozen on dry ice. Lysis was performed in 125 μl ice-cold lysis buffer (50 mM Tris-HCl, pH7.5, 150 mM NaCl, 1 mM MgCl_2_, 0.5% Triton X-100, 1 mM dithiothreitol (DTT), 10% glycerol) with inhibitors (complete EDTA free protease inhibitor, 40694200, Roche, and 1 mM PMSF) using a glass bead beater (Cryoprep). The volume was increased by the addition of 275 μl of ice-cold lysis buffer with protease inhibitors, and the extracts were clarified by two successive rounds of centrifugation. Samples were incubated 1h at 4°C with anti-MYC magnetic beads (TA150044, Origene). Beads were washed 5 times with lysis buffer without inhibitors. Hot reducing agent was added during 5min to eluate proteins.

### Northern blotting

Northern blot experiments were essentially performed as described in (47). RNA was prepared as described in (23). 8□μg of RNA was resolved on a 1.2% agarose gel containing 6.7% formaldehyde in MOPS buffer. After capillary transfer in 10× SSC onto a Hybond N+ membrane (GE Healthcare), RNA was UV-crosslinked and stained with methylene blue to visualise ribosomal RNA bands. For snoRNAs analysis, 16µg of RNA was resolved in 8% Urea-PAGE. Electro-transfer was performed in TBE overnight onto a Hybond N+ membrane (GE Healthcare, RPN303B). Gene-specific probes were generated by random priming in the presence of ATP [α^32^P] using the Prime-It II Random Primer Labeling Kit (Agilent, 300385) using PCR generated DNA template produced from gDNA isolated from a wild type *S*.*pombe* strain (YP71) using oligonucleotides listed in Supplementary Table 8. Probes were added to the membrane and hybridized at 42°C overnight. After repeated washes in 2× SSC, 0.1% SDS, blots were exposed with Amersham Hyperfilm MP (GE Healthcare, 28-9068-44).

### Poly(A)+ RNA interactome capture

Poly(A)+ *S. pombe* RNA interactome capture (RIC) was obtained from (56, 78). Briefly, two sets of triplicate experiments (Wild-type 1 (WT1) + *mtl1-1* = First eXperiment (FX); WT2 + *rrp6*Δ + *dis3-54* = Second eXperiment (SX)) were performed. *S. pombe* cells were grown at 30°C, in Edinburgh minimal media supplemented with glutamic acid (EMMG) with limited amounts of uracil (10 mg/l), and labelled with a final concentration of 1 mg/l 4-thiouridine for 4.5 h. Cells were harvested by filtration, snap-frozen in liquid nitrogen after UV-crosslinking at 3J/cm^2^ in 50 ml PBS and lysed by grinding in liquid nitrogen. Clear lysates were resuspended in oligo-d(T) lysis buffer (20 mM Tris–HCl pH 7.5, 500 mM LiCl, 0.5% Lithium Dodecyl Sulfate (LiDS), 1 mM EDTA; 5 mM DTT, protease inhibitor cocktail IV (fungal) 1:10,000; 5 mM DTT). 1/48^th^ of total volume of whole cell extract (WCE) was used for proteomics analysis, the rest was subjected to pull-down with oligo-d(T)x25 magnetics beads (NEB-S1419S), 1ml of slurry per 1L of cell culture during 1 hour at 4°C. Immobilized oligo-d(T)x25 magnetics beads were washed two times with oligo-d(T) wash buffer 1 (20 mM Tris–HCl pH 7.5, 500 mM LiCl, 0.1% LiDS, 1 mM EDTA, 5 mM DTT) at 4°C, two times with oligo-d(T) wash buffer 2 (20 mM Tris–HCl, pH 7.5, 500 mM LiCl, 1 mM EDTA) at room temperature and two times with oligo-d(T) low salt buffer (20 mM Tris–HCl pH 7.5, 200 mM LiCl, 1 mM EDTA) at room temperature. RNA-proteins complexes were eluted from beads with oligo-d(T) elution buffer (20 mM Tris–HCl pH 7.5,1 mM EDTA), 330µl/L of culture during 10min at 55°C. 1/33 of the oligo-d(T) pull-down total volume was used for RNA sequencing analysis. The rest was subjected to RNase-A and RNase T1 treatment and subjected to mass spectrometry analyses as described in (56).

### Statistical data analysis

Statistical analysis was performed essentially as described in (40) with the following modifications. To be considered for the analysis, protein was required to be present in at least one of the interactomes with two non-zero values. Raw intensities were log2 transformed, normalised to the same median and analysis was followed by the imputation of missing values using a minimal value approach (MinDet – where each sample is considered independently). Data manipulations, principal component analysis (PCA) and Pearson correlation plots were performed with the DEP package implemented in R (1). Median-normalised data values were used to estimate the log-fold changes between exosome mutants and WT cells, which were further normalised by the whole-cell extract values (WCE-normalisation). To minimise the batch effects control experiments (WT cells) were performed twice in triplicates alongside each of the sets of exosome mutants (First triplicate = FX = WT1 + *mtl1-1*; Second triplicate = SX = WT2 + *rrp6*Δ + *dis3-54*). To test the changes between whole-cell extract normalised (WCE-normalised) proteomes of mutants and WT cells, we used modified scripts from the DEP package. Briefly, this software takes advantage of the Limma package that calculates moderated t-statistics on a linear model fit to the expression data (79). It allows defining custom contrasts (like comparing the difference of differences – as in the case of the WCE-normalised intensities). Proteins with a log2 (WCE-normalised RIC of exosome mutant/ WCE-normalised RIC of WT) > 1 were considered specifically enriched in exosomes mutants. Remaining figures and analyses were performed with custom scripts or ones modified from the DEP package. *S. pombe* GO term annotations and information on individual proteins were retrieved using PomBase (80).

### Poly(A)+ RNA Fluorescence *In Situ* Hybridization (FISH)

Poly(A)+ RNA FISH was done as described in (81, 82), using oligo-d(T)(x20)-alexa488 (Invitrogen 7206906) DNA probe. Briefly, 5×10–1×10^8^ cells were used for one hybridization reaction. Cells from an asynchronously growing culture were fixed by the addition of paraformaldehyde into the culture to a final concentration of 4%. Cell pellet was washed with 1ml of buffer B (1.2□M sorbitol, 100□mM KH_2_PO_4_ at pH□7.5, 4□°C) and immediately after, cells were resuspended in 1ml of spheroplast buffer (1.2□M sorbitol, 100□mM KH_2_PO_4_ at pH 7.5, 20□mM vanadyl ribonuclease complex and 20□μM β-mercaptoethanol) with 1% 100T zymolyase (MP Biomedicals, 083209-CF) and cell wall was digested for 60min. The reaction was stopped by washing with 1 ml of cold buffer B. Cells were incubated for 20□min in 0.01% Triton-X100/1X PBS and washed with 10% formamide/2× SSC at room temperature. Before hybridization, 50□ng of the oligo-d(T) probe was mixed with 2μl of a 1:1 mixture between yeast transfer RNA (10□mg/ml, Life Technologies, AM7119) and salmon-sperm DNA (10□mg/ml, Life Technologies, 15632-011) and the mixture was dried in a vacuum concentrator. Hybridization buffer F (20% formamide, 10□mM NaHPO_4_ at pH□7.0; 50□μl per hybridization) was added, and the probe/buffer F solution was incubated for 3□min at 95□°C. Buffer H (4× SSC, 4□mg/ml BSA (acetylated) and 20□mM vanadyl ribonuclease complex; 50□μl per hybridization) was added in a 1:1 ratio to the probe/buffer F solution. Cells were resuspended in the mixture and incubated overnight at 37□°C. After three washing steps (10% formamide/2× SSC; 0.1% Triton-X100/2x SSC and 1x PBS), cells were resuspended in 1× PBS/DAPI and mounted into glass slides for imaging. *Z*-planes spaced by 0.2□μm were acquired on a Ultraview spinning-disc confocal. Acquisition was done with DAPI filter (405nm) and a FITC filter (488nm for alexa488 acquisition). Images were analysed using ImageJ software (83). Signal intensity was measured in the same square surface containing the nucleus (i.e. DAPI) or the cytoplasm. Average intensity of each square was calculated and the ration of Nucleus Signal/ Cytoplasm Signal was calculated for each cell. The mean and the confidence interval of the mean were calculated with α=0.05. Statistical analysis (un-paired *t-test*) was performed using GraphPad Software.

### RNA sequencing

For spike-in normalisation, the *S. cerevisiae* cells were added to *S. pombe* at 1:10 ratio prior to RNA isolation. Total RNA was extracted from cultures in mid-log phase using a standard hot phenol method and treated with RNase-free DNase RQ1 (Promega, M6101) to remove any DNA contamination. For total RNA sequencing, experiments were done in duplicates, and ribodepletion was performed using ribo-minus transcriptome isolation kit (Invitrogen, K155003). Poly(A)+ RNA sequencing was performed by using 1/33 of the oligo-d(T) pull-down total volume, subjected to proteinase K treatment for 1h at 50°C. Poly(A)+ RNA was recovered by a standard hot phenol method. Experiments were done in triplicate. cDNA libraries were prepared using NEBNext® Ultra™ II Directional RNA Library Prep Kit for Illumina (NEB#E7760S) for 50ng of total RNA and using the NEBNext Ultra Directional RNA Library Prep Kit for Illumina (NEB #E7420) for 100ng of WT1, *mtl1-1, rrp6Δ* and *dis3-54* purified oligo-d(T) RNA. Paired-end sequencing was carried out on the Illumina HiSeq 500 platform. RNA sequencing data are available with the GEO number GSE148799 and GSE149187.

### RNA-sequencing data analyses

Quality trimming of sequenced reads was performed using Trimmomatic (Galaxy Version 0.32.3, RRID:SCR_011848). Reads were aligned to the concatenated *S. pombe* (ASM294v2.19) using Bowtie 2 (TopHat) (84). For spike-in normalisation, reads derived from different *S. pombe* and *S. cerevisiae* chromosomes were separated. Reads mapped only once were obtained by SAMTools (85) and reads were mapped to the genome using genome annotation from (86). Differential expression analyses were performed using DESeq2 (87) in R and using the spike-in normalisation. For poly(A)+ RNA sequencing total read count normalisation using DEseq2 (87) in R was used. The significance of RNAs list overlaps was calculated using a standard Fisher’s exact test. For gene ontology analysis, up or down-regulated protein-coding gene lists were submitted to AnGeLi (http://bahlerweb.cs.ucl.ac.uk/cgi-bin/GLA/GLA_input), a web based tool (66).

### ACCESSION NUMBERS

Raw (fastq) and processed sequencing data (bedgraph) can be downloaded from the NCBI Gene Expression with the GEO number GSE148799 and GSE149187. Mass spectrometry data are available via ProteomeXchange with identifier PXD016741.

## ACKNOWLEDGEMENT

We thank the National Bio Resource Yeast Project, S. Grewal, P. Bernard and JP. Javerzat for strains and constructs. We thank members of the Vasilieva’s laboratory and Jean-Paul Javerzat for their valuable comments on the manuscript. We acknowledge the Micron Advanced Bioimaging Unit (supported by the Wellcome Strategic Awards 091911/B/10/Z and 107457/Z/15/Z) for their support and assistance in performing FISH experiments described in this manuscript.

## FUNDING

This work was supported by the Wellcome Trust Senior Research fellowships [WT106994MA to L.V.] and a Medical Research Council Career Development Award [MR/L019434/1 to A.C.].

## CONFLICT OF INTEREST

None declared

## FIGURE LEGENDS

**Figure EV1:**
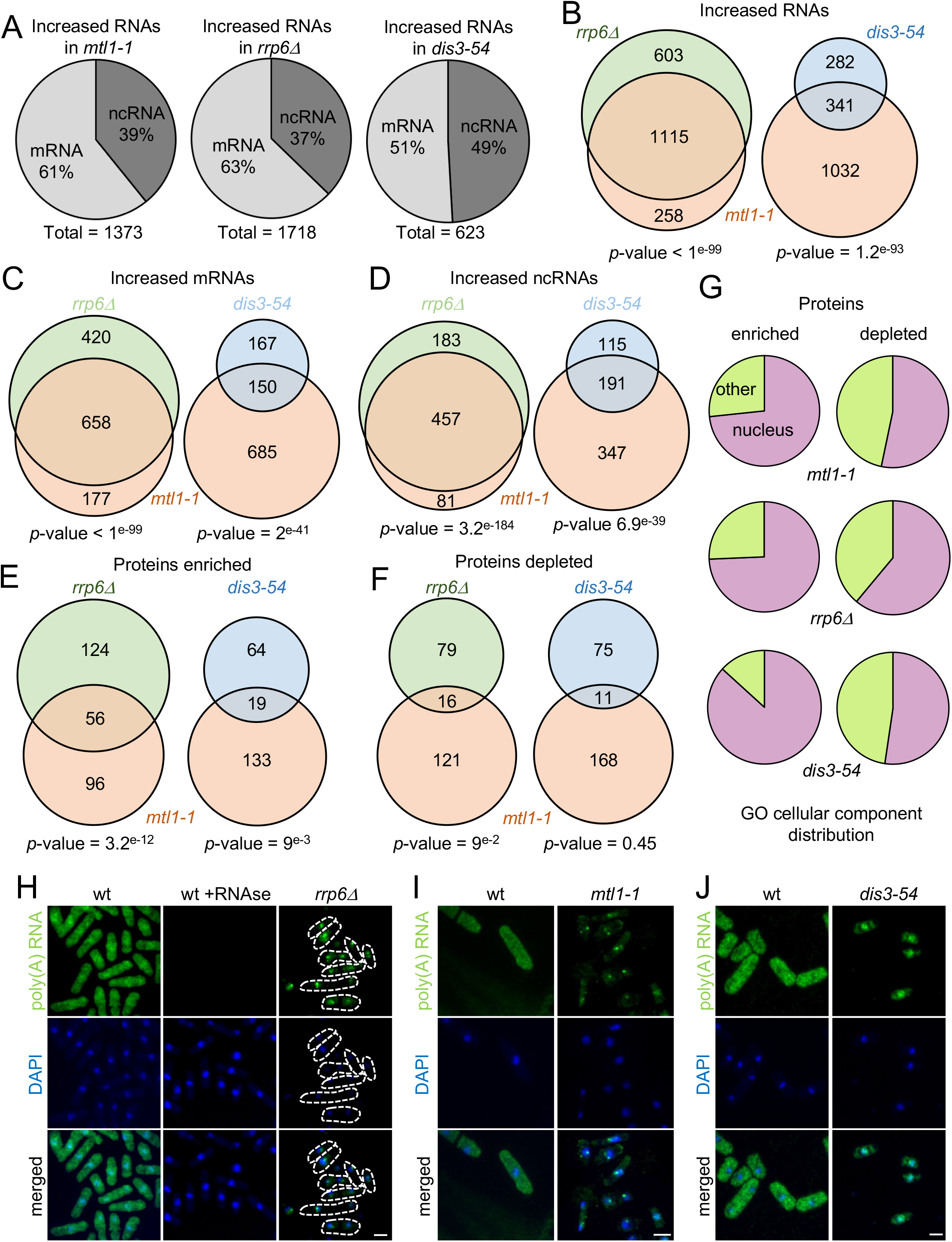
Effect of the exosome mutations on poly(A)+ RNA transcriptome and protein interactome. **A**. mRNAs and ncRNAs are up-regulated (>1.5-fold, *p*-value<0.05) in *mtl1-1, rrp6Δ* and *dis3-54* mutants compared to WT. **B**. Venn diagram showing overlap between RNAs that show increased levels (>1.5-fold, *p*-value<0.05) in *mtl1-1, rrp6Δ* and *dis3-54*. The *p*-values indicate the probabilities that the observed overlaps occurred by chance (6949 genes analysed). **C**. Venn diagram showing overlap between mRNAs increased levels in *mtl1-1, rrp6Δ* and *dis3-54* mutants compared to WT (>1.5-fold, *p*-value<0.05). The *p*-values indicate the probabilities that the observed overlaps occurred by chance (5153 mRNAs analysed). **D**. Venn diagram showing overlap between ncRNAs that show increased levels in *mtl1-1, rrp6Δ* and *dis3-54* mutants compared to WT (>1.5-fold, *p*-value<0.05). The *p*-values indicate the probabilities that the observed overlaps occurred by chance (1796 ncRNAs analysed). **E**. Venn diagram showing proteins commonly enriched in poly(A)+ RNA pull-down of *mtl1-1* and either *rrp6Δ* or *dis3-54* relative to WT. The *p*-values indicate the probabilities that the observed overlaps occurred by chance (1146 proteins analysed). **F**. Venn diagram showing proteins depleted in poly(A)+ pull-down of *mtl1-1* and either *rrp6Δ* or *dis3-54* relative to WT. The *p*-values indicate the probabilities that the observed overlaps occurred by chance (1146 proteins analysed). **G**. RNA association of nuclear proteins (GO:0005575) is primarily affected in the exosome mutants. **H-J**. Poly(A)+ RNA FISH illustrating poly(A)+ RNA accumulation in the nucleus in *rrp6Δ* (**H**) *mtl1-1* (**I**) and *dis3-54* (**J**). DAPI is shown in blue. Poly(A)+ RNA is represented in green. Scale bar = 5µm.

**Figure EV2:**
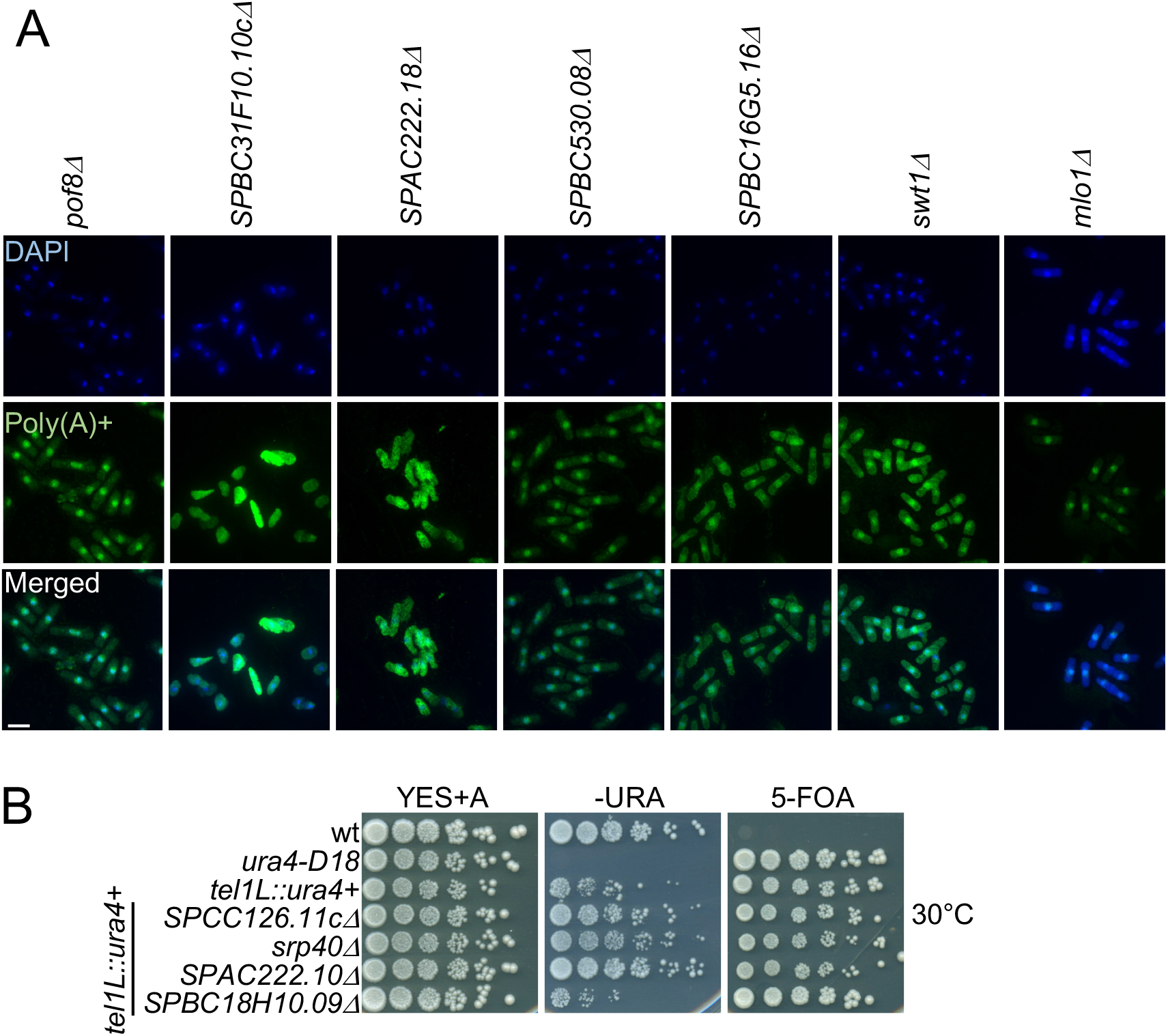
**A**. Poly(A)+ RNA FISH in the indicated strains. DAPI staining is shown in blue. Poly(A)+ RNA is visualised in green. Scale bar = 10µm. **B**. Serial dilutions of the indicated strains plated on complete medium (YES+A), lacking uracil (-URA) and containing 5-FOA. Cells are grown at 30°C. Related to Figure 2E.

**Figure EV3.**
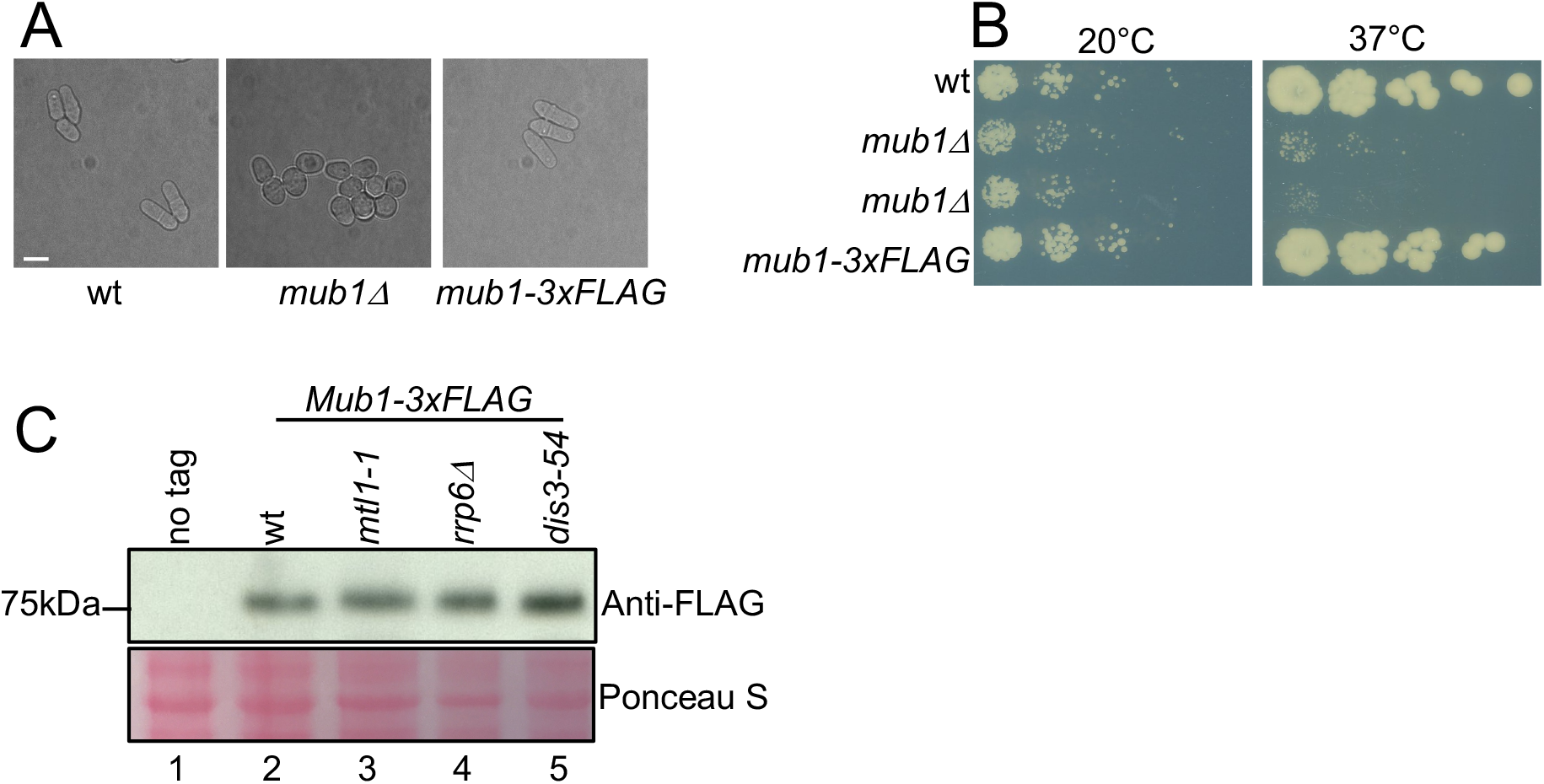
**A**. Mub1 deletion leads to altered cellular morphology. Phase contrast images of the cells from the indicated strains. Scale bar = 10µm. **B**. Serial dilutions of the indicated strains plated on complete medium (YES+A) at the indicated temperatures. **C**. Western blot showing levels of endogenously expressed Mub1-3xFLAG in WT and exosome mutants. Cellular proteins visualised with ponceau S are used to control for loading.

## Notes

### Competing Interest Statement

The authors have declared no competing interest.

### Summary of Updates

the new version is more focused compared to the previous one.

